# Learning Active Multimodal Subspaces in the Brain

**DOI:** 10.1101/2021.12.06.471396

**Authors:** Ishaan Batta, Anees Abrol, Zening Fu, Vince D. Calhoun

## Abstract

Here we introduce a multimodal framework to identify subspaces in the human brain that are defined by collective changes in structural and functional measures and are actively linked to demographic, biological and cognitive indicators in a population. We determine the multimodal subspaces using principles of active subspace learning (ASL) and demonstrate its application on a sample learning task (biological ageing) on a Schizophrenia dataset. The proposed multimodal ASL method successfully identifies latent brain representations as subsets of brain regions and connections forming co-varying subspaces in association with biological age. We show that Schizophrenia is characterized by different subspace patterns compared to those in a cognitively normal brain. The multimodal features generated by projecting structural and functional MRI components onto these active subspaces perform better than several PCA-based transformations and equally well when compared to non-transformed features on the studied learning task. In essence, the proposed method successfully learns active brain subspaces associated with a specific brain condition but inferred from the brain imaging data along with the biological/cognitive traits of interest.

## 1. INTRODUCTION

Understanding structural and functional changes in the brain has been of prime importance in neuroscience. Numerous learning-based approaches have addressed it using neuroimaging data by performing predictive or diagnostic analysis [1], as well as working to identify indicators for brain disorders [2, 3]. Such indicators could be utilized along with certain biological or cognitive traits to create biomarkers based on neuroimaging data. Apart from diagnostic advantage, such approaches may also give insights towards understanding the organization and functioning of the brain. More recently, studies have shown that instead of tracking changes in individual brain areas, mental health disorders are characterized by collectively changing structural as well as functional subspaces in the brain [4]. Many studies have aimed to achieve this by synthesizing multimodal features from neuroimaging data [5, 6]. Thus, it is becoming increasingly emphatic to have extensive frameworks that take into account multiple modalities of brain as well as involve a synthesis as to understand the subspace properties within as well as across modalities at the same time.

Motivated by that, we develop and introduce a multimodal framework based on active subspace learning (ASL) which helps in identification of subspaces from the brain that are defined by collective changes in both structural as well as functional measures. Our framework is based on eigen-decomposition of the covariance of the gradient of a function defined from multimodal input features to a given cognitive or biological target variable. It uses the most contributing eigenvectors to identify the directions (subspaces) in which certain structural components and functional connections co-vary the most in association with the target variable. We show by repeated analysis that the method is stable and is capable of extracting structural components and functional connections which consistently contribute to active subspaces. Thus, our framework is able to: (a) synthesize multimodal features to identify active and sparse subspaces defined by both structural and functional modalities, (b) utilize information from the target variable at hand while computing subspaces rather than generic subspace computation, (c) identify subspace patterns in Schizophrenia that are different from healthy controls and (d) retain predictive information in the transformed features generated by projecting the input data onto active subspaces.

## 2. METHODS

### 2.1. Dataset, Pre-Processing and scICA Components

Structural MRI (sMRI) and resting-state functional MRI (fMRI) data were used from the Function Biomedical Informatics Research Network (fBIRN) [7] dataset. FBIRN dataset includes 160 healthy control (HC) subjects (45/115 F/M, age range 19 −59 yrs, mean age 37.04 ± 10.86) and 151 subjects with Schizophrenia (SZ) (36/115 F/M, age range 18− 62 yrs, mean age 38.77 ± 11.63). Data pre-processing was done using the statistical parametric mapping (SPM12, http://www.fil.ion.ucl.ac.uk/spm/) toolkit and Matlab 2016 followed by registration onto the standard Montreal Neurological Institute (MNI) space and voxel-level gray matter volume (GMV) maps were created from the structural data and fMRI time series from the functional data.

Subsequently, spatially constrained group independent component analysis (scICA) approach [8] with the Neuro-mark framework [9] was used on the functional data to get 53 consistent component masks corresponding to brain areas. Static functional connectivity (SFNC) features were computed using the fMRI time-series from the 53 components, giving ^53^*C*_2_ = 1378 pairwise correlations for each subject. The masks were registered to the structural space to obtain 53 structural features corresponding to mean scICA loadings from voxel-level GMV maps.

### 2.2. Active Subspace Learning Framework

For a given point **x** ∈ ℝ^*m*^ in an input feature space of dimension *m*, let *f* : ℝ^*m*^→ ℝ be a function mapping the space of input features to the space of target variable. Note that in our case, *x* could be considered as features from the brain, *y* as a cognitive or biological trait, and *f* could be a regression function. The methodology of Active Subspace Learning Analysis [10] (ASL) works by performing the eigen-decomposition of the expected outer-product (covariance) of the gradient of function *f* as follows:

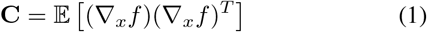

In practice, one can estimate **C** as **Ĉ** from the data. In this approach, we considered *f* to be the Gaussian Process Regression function, as a closed-form estimation of the matrix **Ĉ** can be obtained from a dataset of *n* subjects, [**X, y**] with **X** ∈ ℝ^*m×n*^, and **y** ∈ ℝ^*n*^ [11].

The eigen-decomposition of **C** is then utilized to identify a set of active subspaces defined by the eigenvectors with significantly large eigenvalues. Further, one can create a set of transformed features 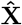 by projecting the input space onto these active subspaces.

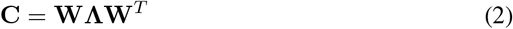

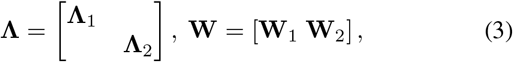

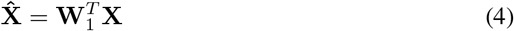

Essentially, the active subspace vectors are the directions in which the entity represented by the features shows the most variation in association with the target variable at hand.

### 2.3. Identifying Multimodal Subspaces for the Brain

For neuroimaging data, this framework can be used towards a dual purpose of identifying subspaces and generating transformed features for learning applications or subsequent analysis. In the case of structural features as input, each column of the **W**_**1**_ matrix represents an active subspace made up of a subset of structural components of the brain, where the value corresponding to each component signifies the contribution of the component in the particular subspace. For SFNC features as input, the active subspaces would correspond to a subset of connections (sub-networks) in the brain. In this work, we concatenated the structural and SFNC inputs to create a multimodal framework that is used to obtain subspaces spanning both structural and functional domains i.e. the input matrix **X** now can be written as 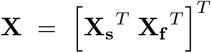, where **X**_**s**_ and **X**_**f**_ are matrices for structural and SFNC features respectively. The active subspaces thus obtained correspond to the direction of highest change with respect to the target trait in a multimodal dimension made of structural and functional subdimensions. We selected age as the target variable.

## 3. RESULTS

### 3.1. Active Multimodal Subspaces

Multimodal ASL analysis was done as described in subsections 2.3, 2.2. The eigen-matrix was computed as **W** = [**W**_1_ **W**_2_], where **W**_1_ represents the multimodal active subsapce matrix separated from **W**_2_ based on a threshold on the eigenvalues cumulatively capturing 99% of the variance. 100 most predictive SFNC features were used for computational feasibility of eigen-decompositions involved due to repetition experiments discussed subsequently.

It can be observed that the multimodal subspaces are indeed sparse (Figure 1a) with each mulimodal subspace consisting of contributions from both structural components as well as functional connections. Further, to check how the active subspaces vary between control (HC) and Schizophrenia (SZ) groups, we computed active subspaces using data from only HC and only SZ groups. Figure 1b shows the group differences between means of contributions to active subspaces i.e. elements of the **W** matrix in Equation 3 between HC and SZ groups for 100 reps. It can be noted that many components/connections that define the subspaces in Schizophrenia are significantly different from the ones for the control group.

**Fig. 1:**
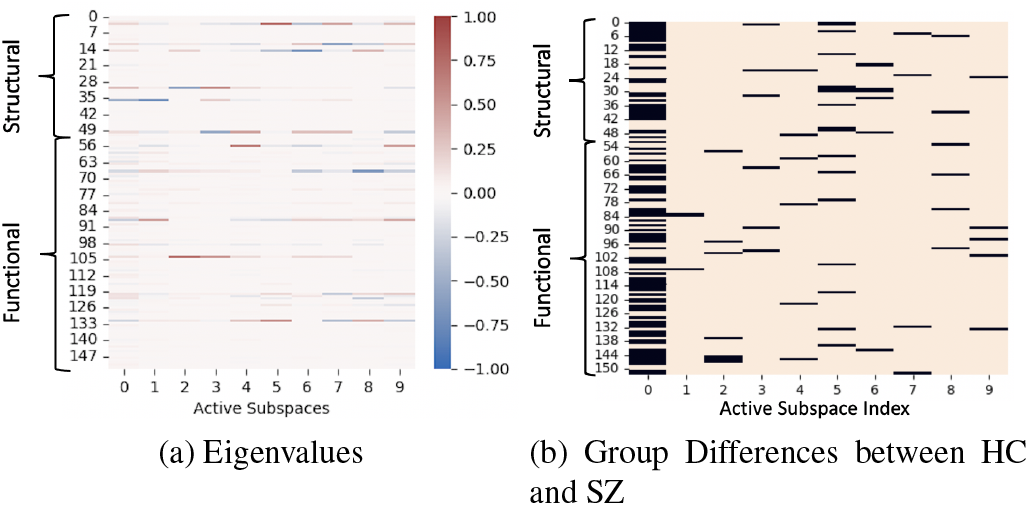
(a) Matrix representing active multimodal subspaces for a single run of the ASL framework. Each column is an active subspace, while each row corresponds to the contribution of the structural component/functional connection in the active multimodal subspace. (b) Group differences between active subspaces computed using data from only control (HC) and only Szhizophrenia (SZ) groups. Each element at position *i, j* corresponds to the two-sided t-test result on whether the mean value of corresponding element in subspace matrix (i.e. **W**_*ij*_) across 100 repetitions is significantly different (*p <* 0.05) between the HC and SZ groups. Elements with significant difference are shown in black.

### 3.2. Subspace Stability and Consistent Components

To check the robustness of the subspaces, the whole ASL analysis was performed for 100 repetitions on randomly selected 80% of the data as training set. To test for any structural components or functional connections that feature consistently in active multimodal subspaces across the repetitions, we defined a contribution metric vector **s**. For the *r*^th^ repetition, the *c*^th^ element of **s**^(*r*)^ is defined for the corresponding component/connection *c* as,

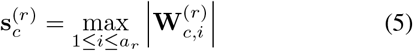

where *a*_*r*_ is the number of active subspaces and **W**^(*r*)^ is the subspace matrix for repetition *r*, as defined in Equation 3. 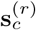 essentially captures the extent to which the *c*^th^ component/connection contributes to any of the active subspaces for the *r*^th^ repetition. Since each subspace vector is a unit basis vector of an eigen-space, the maximum value across subspaces in Equation 5 accounts for the significant contribution in at least one active subspace and the absolute value accounts for weight in any direction. The summarized contribution 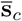 of component/connection *c* was computed by simply taking the mean across repetitions. For each component, a one-sided t-test was done on the values 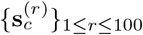 to check if the mean 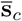 was significantly high 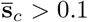 with *p -*value *<* 0.05). Based on this test for consistent contribution of components/connections, 9 structural components and 27 functional connections were found to significantly contribute to the multimodal subspaces (See Figure 2).

**Fig. 2:**
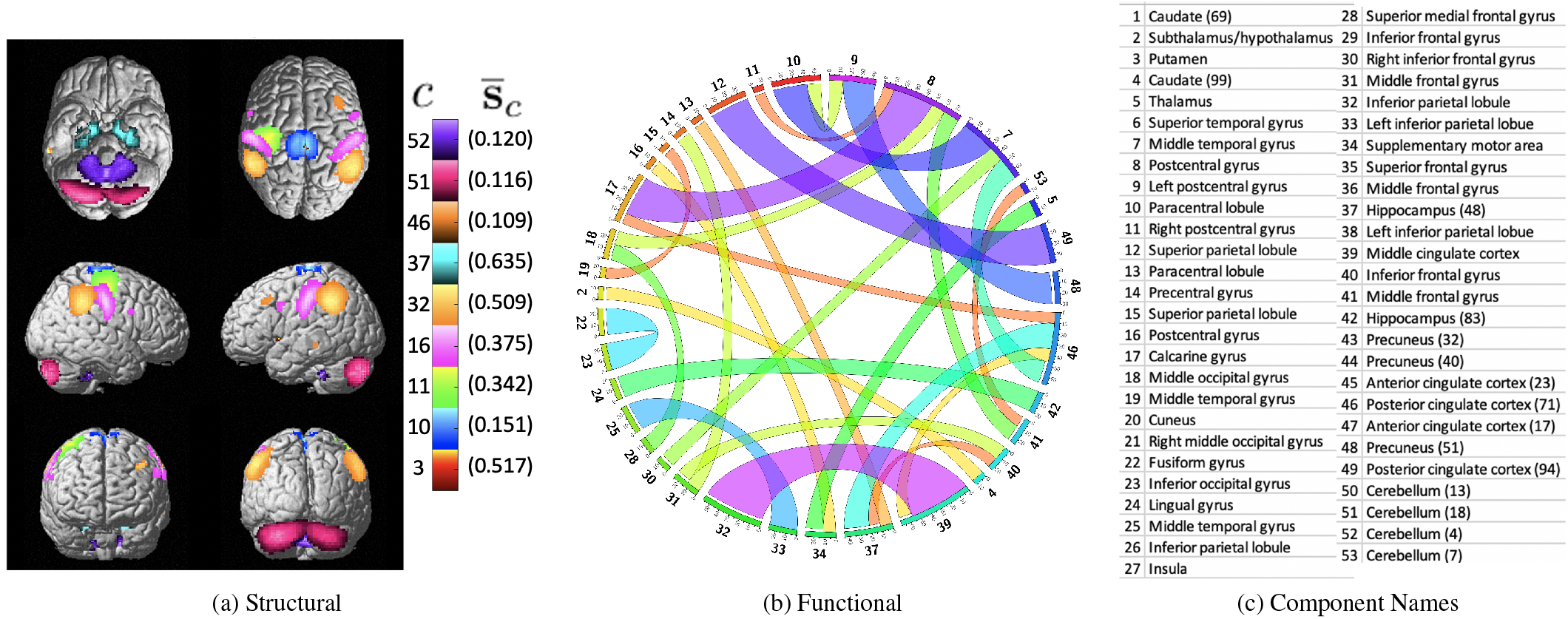
Components/connections with significantly high value of mean overall contribution 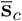 as defined in subsection 3.2. (a) Significantly contributing structural components are shown on the brain map along with 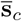 values and (b) Functional connections are shown between components as connectogram with widths representing the value of 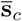 The numbers on the left of the colorbar in (a) and outside the circumference of the connectogram in (b) correspond to the component names in (c).

### 3.3. Performance Comparison

Regression analysis was also done on the ASL transformed features 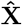 in Equation 4 to check if they retain or enhance predictive information. Figure 3 shows performances from 100 repetitions of Ridge regression analysis with 4-fold cross validation on the training set and testing on randomly chosen 20% held-out data. Performance of ASL features created as in Equation 4 was compared with baseline models including null model (predicting mean of the training data), non-transformed multimodal features (Raw), and finally principle component analysis (PCA)-based transformation methods which do not take the target variable information into account. These include standard PCA, kernel PCA with linear (kPCAl), radial kernels (kPCAr), and sparse PCA (sPCA).

**Fig. 3:**
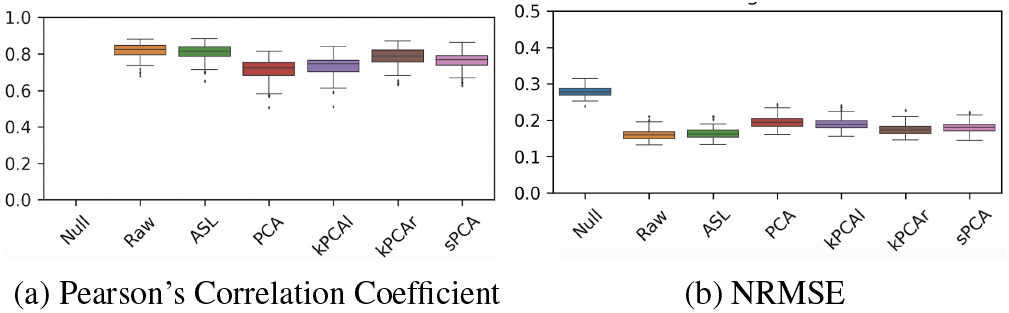
Performance comparison for regression analysis on age using Ridge regression. (a) Pearson correlation and (b) normalized root mean square error (NRMSE) are plotted for ASL features computed by projecting the concatenated multimodal features onto the active multimodal subspaces as computed in Equation 4. Comparison is shown with null model, non-transformed multimodal features (Raw) and PCA-based transformations described in subsection subsection 3.3. (Note that null model has no correlation as the prediction is a constant vector)

ASL feature performance was significantly better than the PCA based methods and comparable to non-transformed high dimension multimodal features, indicating that the predictive information is retained while computing the multimodal subspaces and projecting input data as in (Equation 4).

## 4. CONCLUSION

Our framework is able to identify sparse, stable subspaces which synthesize both structure and function, while taking into account the association with changes in a given target variable (age), which is also a possible reason for retention in predictive performance Figure 3. As depicted in Figure 1b, Schizophrenia involves significantly different brain subspace patterns than the controls in characterizing the changes in the target variable. As in Figure 2, it can be seen that there are multiple brain regions whose structure as well as function co-vary in association with the target variable, while for certain regions only one of the modalities is involved. Rather than looking at individual modality or individual regions, the framework presented in this work takes into account subspaces that span both functional and structural aspects of the brain and is also able to compute subspcaces specific to a given disorder. Additionally, having a target variable into consideration makes it viable to identify multimodal biomarkers for disorders involving a particular cognitive or biological trait of interest. In future this work could be extended to create more robust analysis scalable to higher dimensions as well as faster in computation.

## 5. ACKNOWLEDGMENTS

Research reported in this publication was supported by National Institute of Mental Health (HHS - NIH) of the National Institutes of Health under award number R01MH118695.

## Notes

### Competing Interest Statement

The authors have declared no competing interest.

